# Brain size reduction in dogs was already established at least by the Late Neolithic of western Europe, 5,000 years ago

**DOI:** 10.64898/2025.12.12.693870

**Authors:** Thomas Cucchi, Lucile-Morgane Hays, Alessio Veneziano, Margot Michaud, Colline Brassard, Rose-Marie Arbogast, Pierre Pétrequin, Mietje B. Germonpré, Evelyne Crégut-Bonnoure, Frederic Elleboudt, Kalman Czeibert, László Zsolt Garamszegi, Enikő Kubinyi, Niclas Kolm, Tibor Csörgő, Justine Joseph, Salomé Leroy, Claude Guintard, Marion Fusellier, Christophe Duchamp, Anthony Herrel, Loukas George Koungoulos, Thomas J. Peachey, Lachie Scarsbrook, Laurent Frantz, Joan Madurell-Malapeira, Sandrine Ladevèze

## Abstract

The timing and causes of brain size reduction in domestic dogs remain uncertain. Using endocast’s volume as a proxy for brain size, this study provides a first insight into long-term brain size evolution in the wolf-dog lineage. We compared endocranial volumes of 185 modern and 22 prehistoric wolves and dogs ranging from Western Europe to Australia, and spanning the Pleniglacial (35 Ky BP) to the Late Neolithic (5 Ky BP). Our results reveal that Pleistocene so called “protodogs” show no brain size reduction compared to coeval Pleistocene wolves. Instead, we observed a slightly larger relative endocranial volume in the 35,000-year-old ‘protodog’ from Goyet, which could suggest increased behavioural flexibility in the presence of humans. This hypothesis needs to be tested further. In contrast, Late Neolithic dogs show a drastic 46% brain size reduction with an endocranial volumes comparable to modern small terrier and toy breeds. The anxious and wary temperaments of these Late Neolithic dogs, induced by the brain tissue reorganization associated with such a size reduction, could have served an alerting purpose, among the many other potential roles dogs could have played within this Late Neolithic socio-ecosystems.

## Introduction

Reduction in brain size is one of the most widely reported biological consequences of domestication (Balcarcel et al., 2022), often cited as the most reliable marker of the domestication syndrome (Wright et al., 2020). Proposed explanations for this decrease include relaxed selection from predation, foraging, and mating demands, associated with the high energetic costs of the maintenance of neural tissue (Kruska, 1988), reduced cognitive challenges in the captive environment (Yamaguchi et al., 2009), and behavioural selection (Kruska, 2005), with the pleiotropic effects of selection for tameness (Wilkins et al., 2014). Yet, its evolution in the course of the multi millennial domestication history, is still unknown (Hecht et al., 2023).

Among the domesticated mammals, dogs have had the most important brain size reduction, with an average decrease estimated at 20-30% (Balcarcel et al., 2024; Garamszegi et al., 2023). So far, the current understanding of the dog brain size evolution would assume a two-phase trajectory: an initial reduction during the wolf-to-dog transition (Hecht et al., 2023), followed by later increases in specific brain regions under selection for more complex tasks in a more challenging human environment (Kruska, 2005). However, this undated theory has been made from modern comparison, while recent archaeological and paleogenetic studies indicate that dog domestication occurred at least from 20,000 years ago, since they accompanied the first humans dispersal to the Americas (Ní Leathlobhair et al., 2018). Other zooarchaeological studies have even suggested that the domestication process could have occurred as early as 35,000 years ago, with morphometrically divergent specimens described as “protodogs” (Germonpré et al., 2009; Ovodov et al., 2011). Yet, some have argued that this brain size reduction rather reflects very recent breed formation over the last 200 years, when strict aesthetic standards and intensive artificial selection were imposed (Bergström et al., 2022; Lord et al., 2020a). And finally, neuroscientific studies of modern dogs suggest that recent breeding selection has also increased the cerebral cortex to enhance behavioural flexibility and sensitivity to human social cues (Barton et al., 2025).

Therefore, to understand the evolution of dogs’ brain, the archaeological record is required (Bogaard et al., 2021), as well as a modern comparative that covers as much dogs’ behavioural diversity as possible, including populations unaffected by recent selective breeding (Hansen Wheat et al., 2020; Lord et al., 2020b). Such populations exist as free-ranging dogs from around the world, including dingoes and village dogs. Dingoes descend from East Asian dogs that were introduced to Australia at-least 3,300 years ago (Balme et al., 2018; Koungoulos and Fillios, 2020) and have since adapted to become Australia’s apex terrestrial predator, living largely independent of humans (Shipman, 2021). Village dogs, in turn, are free-ranging populations that live and reproduce with little direct human intervention and represent the vast majority of the global dog population (Coppinger and Coppinger, 2001).

In this study, we provide the first insight into dog’s brain size evolution through the analysis of endocast reconstructed from archaeological crania. To do so we assessed endocranial volume (ECV) from Computed Tomography (CT)-derived endocasts as a proxy for brain size (Garamszegi et al., 2023), analysing both absolute ECV and relative ECV (rECV), using cranial measurements to account for body size differences. Because body size can evolve more rapidly than brain size in carnivorans (Michaud et al., 2022), absolute brain size remains an important measure in its own right. Moreover, a recent neuroanatomical study suggests that absolute brain size might bear more information about behaviour and temperament of dogs than relative brain size (Hecht et al., 2021). Finally, to integrate neurocranial morphology with brain size, we also took linear skull measurements from 3D models of the same specimens, and compared them with previously-published metrics from modern and ancient canids. analysing the cranium of modern and ancient canids that predate the onset of intensive selective breeding.

The archaeological dataset include Pleniglacial (> 20,000 BP) and Post-glacial wolves from Belgium, two so-called “protodogs”, namely a Pleniglacial, ~ 35.000 BP specimen from Goyet (Belgium) (Germonpré et al., 2009) and a Upper Palaeolithic, ~ 15,000 BP specimen from Baume Traucade (France) (Germonpré et al., 2025). It also include a rare assemblage of well-preserved Late Neolithic wolves and dogs, from a 5,000, 4,500-year-old lakeshore site (Giligny et al., 1995; Pétrequin, 1997). The modern dataset includes wolves that lived in their natural habitat with a wide diversity of dogs including, dingoes, village dogs and dogs breeds covering all functional behaviours selected over the last 250 years (Serpell, 2017). With this dataset, we assessed (1) whether Upper Pleistocene “protodogs” display neurocranial size differences compared to Pleistocene and Holocene wolves, consistent with an early domestication-linked brain size reduction and (2) whether the transition to the Neolithic farming socio-ecosystem represent an important step in dogs brain size evolution and how these changes compare with those observed in modern breeds and free-living populations (dingoes and village dogs).

### Data collection

We collected CT-scans from 163 crania of adult modern wolves and dogs from six different institutions (details in SI Table 1), along with 22 crania of prehistoric adult wolves and dogs from Belgium and France (details in SI Table 2). We only CT scanned the cranium of specimens having their four permanent upper molars, in order to exclude juvenile specimens. Thus, all specimens were at least seven months old and sexually mature. CT acquisitions were obtained at different facilities and under varying acquisition parameters (details SI Table 3). We only selected specimens that showed no major damage to the neurocranium, which could compromise the virtual reconstruction of the endocast.

The modern wolf dataset (n=59) includes 58 wild males and females from France that were shot during a legal control campaign (2020-2022) in southern France performed by the French Biodiversity Office (OFB) or collected after death (either from natural causes or collisions with cars). We also included one 19^th^-century wolf from the Ardennes region in Belgium. The modern dog dataset (n = 104) includes: (1) 19 village dog skulls from the London Natural History Museum (NHM) that were collected during the 19^th^ and 20^th^ centuries across the world (Russia, Nepal, Chile, Japan, India, Egypt, Sudan, Malaysia, Arabian Peninsula, etc.); (2) 21 dingoes from the Australian Museum collected from the same region of arid inland Western Australia between 1950-1954 and are thought to represent a single population; and (3) 64 dogs selected from the Eötvös Loránd University dog skull collection (Czeibert et al., 2024) that includes 44 breeds across 17 phylogenetic clades (Parker et al. 2017) and seven traditional functions according to the American Kennel Club’s (AKC, www.akc.org). These functional grouping include: (1) The Working includes the oldest and largest breeds, selected to assist humans with various working duties such as pulling carts or sledges, guarding and protecting. (2)The Toy includes the smallest breeds, which are used for companionship. (3) The Herding includes breeds that are known for their trainability and natural intelligence, and that were developed for moving stock (sheep, cattle, reindeer). (4) The Sporting includes breeds used to assist during hunting by locating and retrieving feathered game. (5) The Hound includes breeds selected to chase and pursue warm-blooded quarry. (6) The Non-Sporting includes a wide variety of breeds used as watchdogs and house dogs, mainly sought as companion dogs that are good with people. (7) The Terrier includes breeds selected for hunting, vermin control and guarding.

The archaeological wolf and dog dataset (n = 22; Figure 1, details in SI Table 2) included skeletally mature specimens collected from sites in Belgium and France. Specimens from Belgium include two Pleistocene wolves, one Pleniglacial “protodog” according to skull morphometrics (Germonpré et al., 2009), one Postglacial wolf and one Neolithic wolf from Antwerpen (Hasse, 1909). Specimens from France include a recently excavated and studied Late Glacial “protodog” from France (La Baume Traucade) (Germonpré et al., 2025). The five wolves and ten dogs from the Middle/Late Neolithic lake dwelling site of Chalain were collected among refuse deposits with other animal remains and not as part of a burial (Pétrequin, 1997). Wolves and dogs are easily distinguishable from linear measurements of skulls and long bones (Arbogast et al., 2005). The dog remains of Chalain are mainly represented by crania and mandibles (Chenevoy and Chaix, 1985; Hue, 1910) but also by ornaments made from canines and metapodials (Maréchal et al., 1998), with no obvious butchery marks on the bones, which together suggest a special status for these dogs (Arbogast et al 2005). The crania studied here come from a refuse area along with other animals’ remains. The shoulder height of the Chalain dogs, based on long bone measurements, is about 35 cm (unpublished data), and their skull morphology suggests a distant resemblance to modern herding breeds (Chenevoy and Chaix, 1985).

**Figure 1:**
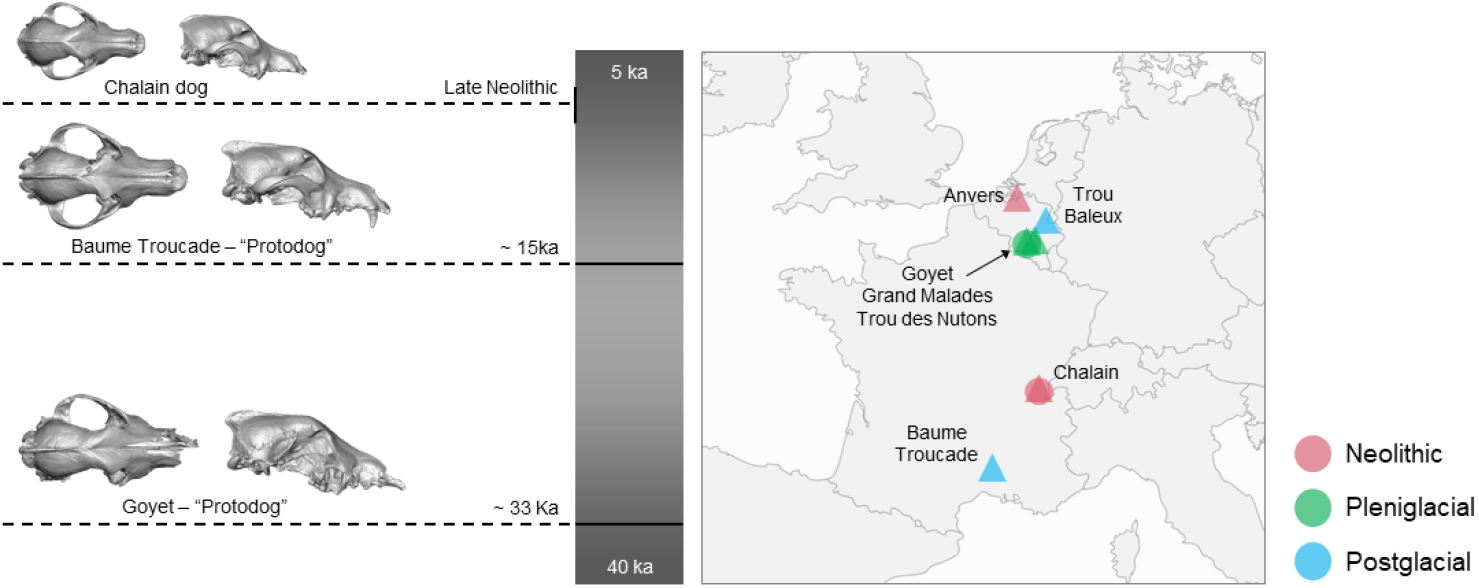
**a)** Timeline with cranium dorsal and lateral views of the Pleniglacial (Goyet) and Late Glacial (Baume Traucade) protodogs and one Late Neolithic dog of Chalain. **b)** Map showing the location of the archaeological sites studied.

### Virtual endocasts and skull measurements

Virtual reconstructions of endocasts (Figure 2) from conventional CT were semi-automated from DICOM files using the “Wrap Solidify” extension for 3D Slicer (freeware, open source, https://www.slicer.org (Andress, 2019)). Their volume in mm^3^ (ECV) was then automatically calculated with the 3D Slicer module. Virtual reconstructions of endocasts from μCT were automated using the AST-3D tool introduced in (Profico et al., 2018). The AST-3D tool works on a 3D cranial mesh intermediate, which we obtained through isosurface interpolation of the μCT volumes using the function *vcgIsosurface* in the R package *Rvcg* (Schlager, 2017).

**Figure 2:**
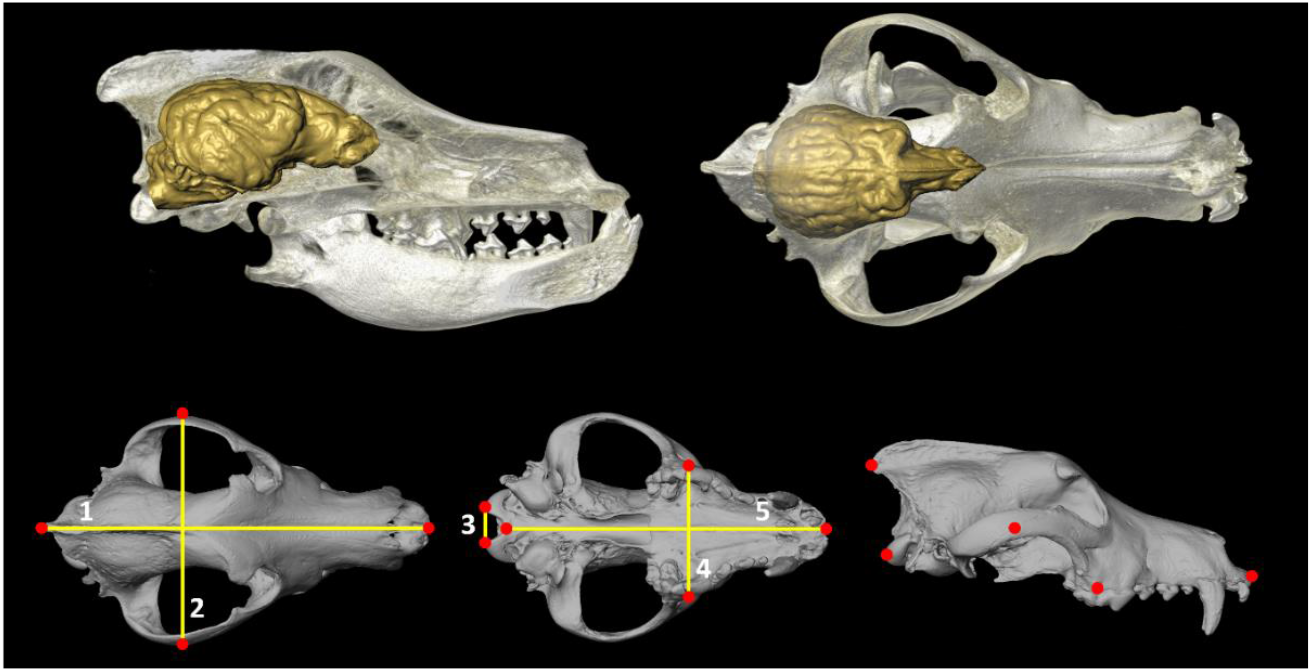
Volume-rendered skull model of a 19^th^-century wolf (RBINS) with the endocast approximating the brain positioned within the braincase, in right lateral view (at the upper right, midsagittal section), and dorsal view (at the upper left). The lower panels from left to right show dorsal, ventral and right lateral views of the wolf cranium with anatomical landmarks for the measurements taken in this study: 1. total cranium length (TL), 2. cranium width, 3. foramen magnum breadth (FMb), 4. greatest palatal breadth (GPB), 5. basal length (BL). Not to scale.

Measurements (Figure 2) were taken from the cranium 3D model using Meshlab (Cignoni et al., 2011). We recorded five measurements: (1) total cranium length (TL), (2) maximum cranium width (W), (3) foramen magnum breadth (FMb), (4) greatest palatal breadth (GPB), and (5) basal cranium length (BL). We then calculated the skull index or cephalic index (CI) as follows: (cranium width / cranium length) *100. According to their CI, domestic dogs where then organised in head groups (dolichocephalic, mesocephalic and brachycephalic) following (Garamszegi et al., 2023). TL and CI of modern dog breeds were compiled from multiple published sources (Balcarcel et al., 2024; Czeibert et al., 2020; Stone et al., 2016). Among the 59 wolves from France, we measured 12 crania. We also collected TL data for Pleistocene, Mesolithic, and Neolithic Eurasian wolves and dogs from the literature (See details in SI Table 4).

### Statistics

Basic statistics (mean, standard deviation, minimum, maximum) were collected for the ECV and TL (SI Table 5). Since variation among the different functional groups of dogs (AKC), wolves and archaeological wolves and dogs were not homogeneous for ECV (Levene’s test: *Df* = 157,18; *F-value* = 2.226 *P* = 0.005) and rECV (Levene’s test: *Df* = 106,14; *F-value* = 2.833 *P* = 0.004), we compared differences among groups using non-parametric Kruskal-Wallis and Dunn’s tests with Bonferroni correction for pairwise comparisons. Differences in variation of TL, ECV and rECV values across groups were visualised with box plots.

To compare ECV and rECV differences across dogs, we grouped them according to their head categories (dolichocephalic, mesocephalic and brachycephalic) following (Garamszegi et al., 2023), and by their AKC traditional functions.

We compared rECV across modern and ancient specimens using linear regression of ECV (dependent) against total cranium length (TL) as the independent variable, including factors distinguishing wolves, dingoes, village dogs and cephalic index categories of dog breeds. To test the difference in rECV variation across modern and ancient specimens, while taking into account the allometric relationship between brain and body sizes, we used the residuals of the linear regression model as rECV values following (Garamszegi et al., 2023).

All statistical analyses were performed with R version 4.4.2. (R Core Team, 2020).

## Results

### 1. Skull size in modern and ancient dogs and wolves

Ancient and modern wolves and dogs display significant differences in skull length (Kruskal-Wallis chi-squared = 160.35, *Df* = 33, *P*< 2.2e-16). Wolves from Pleistocene Belgium (mean=263 mm), Neolithic of France (mean=245 mm) and modern France (mean=239 mm) are in the same cranium size range with what seems a size reduction trend over time (Figure 3) although no significant differences were found among these three groups (SI Table 6). With the exception of one individual from Eliseevichi (256 mm), “protodogs” display shorter crania (mean=229 mm) than broadly contemporaneous Pleistocene wolves, yet no significant differences are observed among the two groups (SI Table 6). The Mesolithic dog from Portugal (Muge) has a shorter cranium than all the wolves, while Mesolithic dogs from Siberia (Zhokhov) are within the lower range of the “protodogs” and modern wolves. The Pleistocene “protodogs” from Belgium (Goyet) and France (La Baume Traucade) are within the lower cranium size range of modern wolves from France. Neolithic dogs from western Europe have shorter skulls than Mesolithic dogs from across Eurasia. and are significantly shorter than dingoes but not than village dogs. Chalain dogs have a similar size range than Terrier dogs but their cranium is not as reduced as those of Toy dogs. Because of the extreme skull size reduction in Toy dogs, we excluded this group from the following analyses of relative ECV across modern and ancient canids.

**Figure 3:**
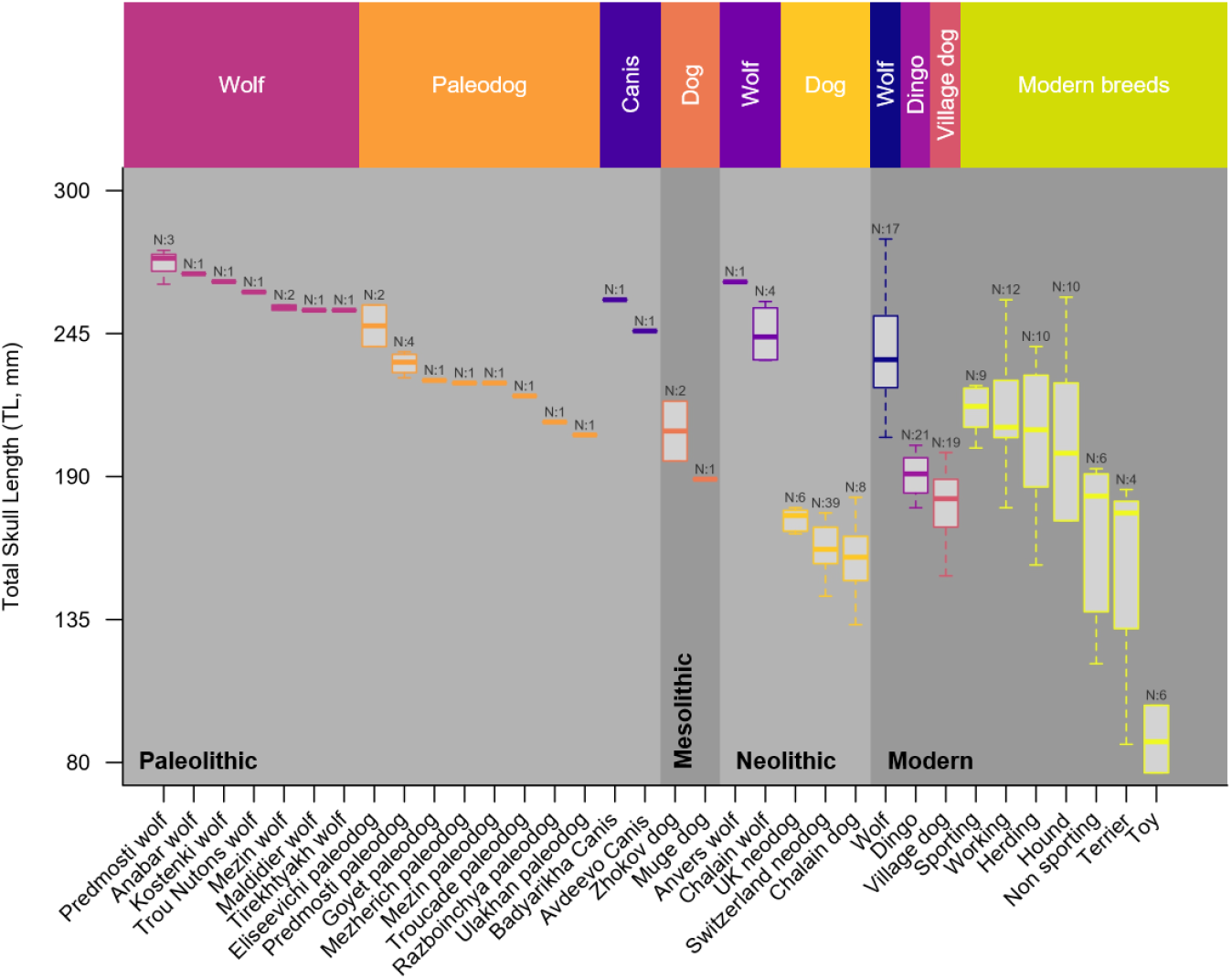
Boxplot of total cranium length (TL) variation in wolves and dogs across Pleistocene, Mesolithic, Neolithic and modern periods. For each group and period, the TL values are displayed in a descending order. TL values include data from this study and from the literature (see SI Table 4). Palaeolithic “*Canis”* from Badyarikha and Avdeevo are specimens without identification as wolf or “protodog”. Dog breeds are grouped according to their AKC classification, which reflects their traditional function.

### 2. ECV variation in modern and ancient dogs and wolves

ECV variation differs greatly across modern and ancient wolves, dingoes and dogs (Figure 4, (Kruskal-Wallis chi-squared = 137.56, *Df* = 14, *P* < 2.2e-16). In this distribution, all dogs (dingoes, village dogs, dog breeds, and Late Neolithic dogs) have significantly smaller ECV than all wolves (Pleistocene, Neolithic and modern) (SI Table 7) with a 32% reduction, which remains the same when we only compare current dogs (dingoes included) and modern wolves. The ECV of wolves across Pleistocene (mean= 153.6 mm^3^), Late Neolithic (mean=140.45 mm^3^) and modern (mean= 133.71 mm^3^) periods range between 111 mm^3^ and 172 mm^3^, with what appears to be a reduction over time. ECV values from Late Neolithic wolves (Chalain) are not significantly different from those of modern wolves (SI Table 7), and are in the range of Pleistocene “protodogs” from Belgium (Goyet) and France (La Baume Traucade).

**Figure 4:**
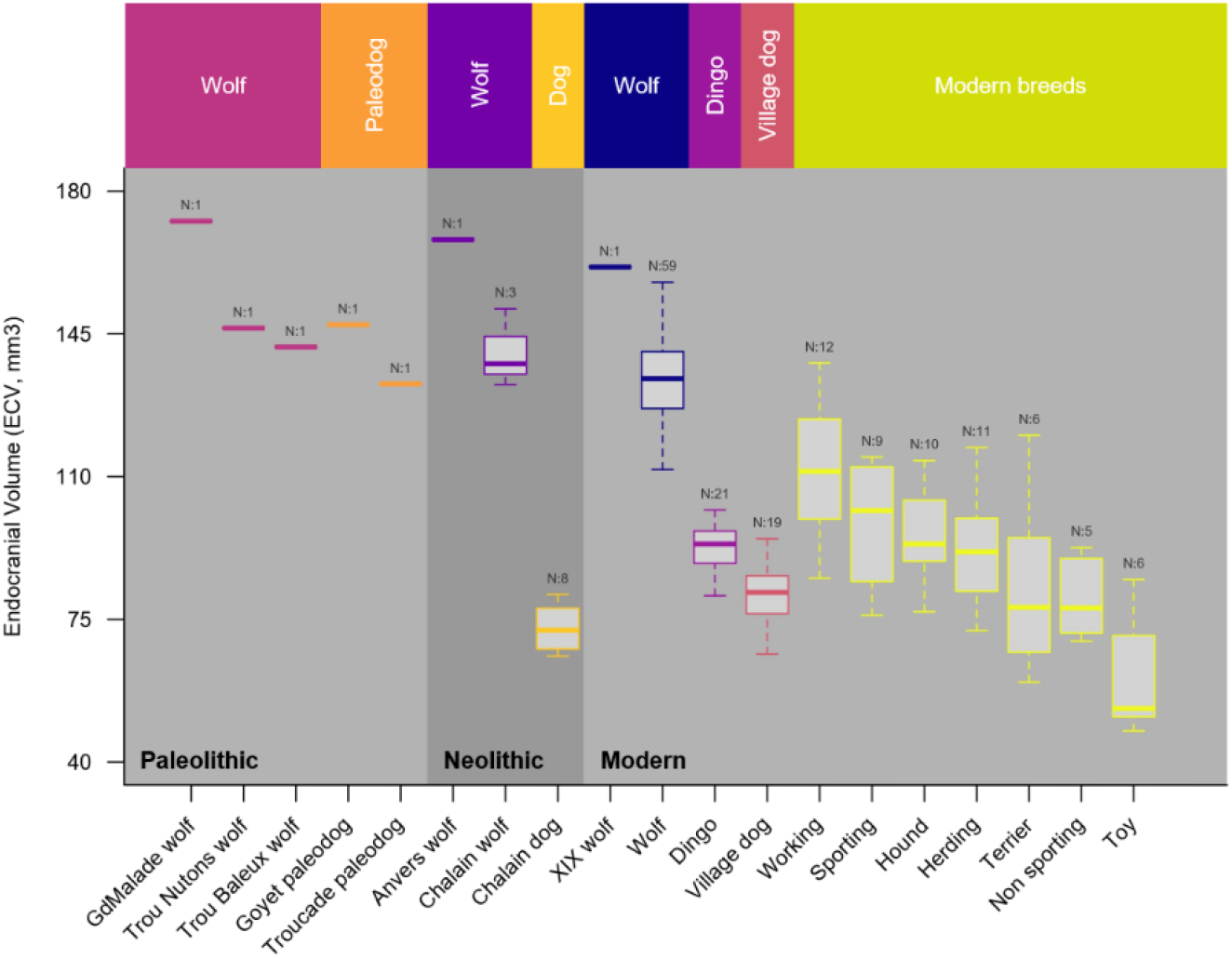
Boxplot of ECV variation across ancient (Paleolithic and Neolithic) and modern wolves, dingoes, and dogs (village dogs and modern breeds). Dog breeds are grouped according to the AKC classification, which reflects their traditional function.

ECV varies greatly among the different dog lineages (i.e., dingoes, and village and breed dogs) (Figure 4) though we only found a significant difference between the largest ECV of working dogs (mean=111.17 mm^3^) and the smallest ECV of Toy dogs (mean=60.14 mm^3^), 17% smaller than all other dog breeds (SI Table 5). Dingoes (mean= 92.47 mm^3^) display significantly larger ECV than village dogs (mean=79.94 mm^3^) but both are in the middle of the dog’s breeds ECV range, between the largest working dog (138 mm^3^) and the smallest Toy dog (47.6 mm^3^). In this wide distribution, the Late Neolithic dogs of Chalain fall between village dogs and Toy dogs’ variation, at the lowest end of the Terrier range with a 46% ECV reduction compared to modern and Neolithic wolves. Seven of the eight Chalain dogs have an ECV similar to modern Medium Spitz, Cocker Spaniel, Pug and Collie, and one has the same ECV as Toy dogs like Chihuahua and Pekingese breeds (SI Figure 1).

### 3. rECV variation in modern and ancient dogs and wolves

Cranium length (TL) explains ~64% of the ECV variation (R^2^ = 0.636, F-statistic = 213.7, *Df* =121, *P* < 2.2e-16) while the foramen magnum breadth (FMb) explains ~52% (R^2^ = 0.518, F-statistic = 75.08, *Df* =68, *P*<0.0001). The variation of rECV across modern wolves and dogs (Figure 5a), shows that wolves have significantly greater rECV than dogs (SI Table 8). The Goyet “protodog” also displays greater rECV than the Pleistocene and Postglacial (Trou des Nutons and Trou Balleux), Late Neolithic, and modern wolves of similar cranium length (Figure 5b). On the other hand, La Baume Traucade Pleistocene “protodog” shows rECV values comparable to those of modern French wolves.

**Figure 5:**
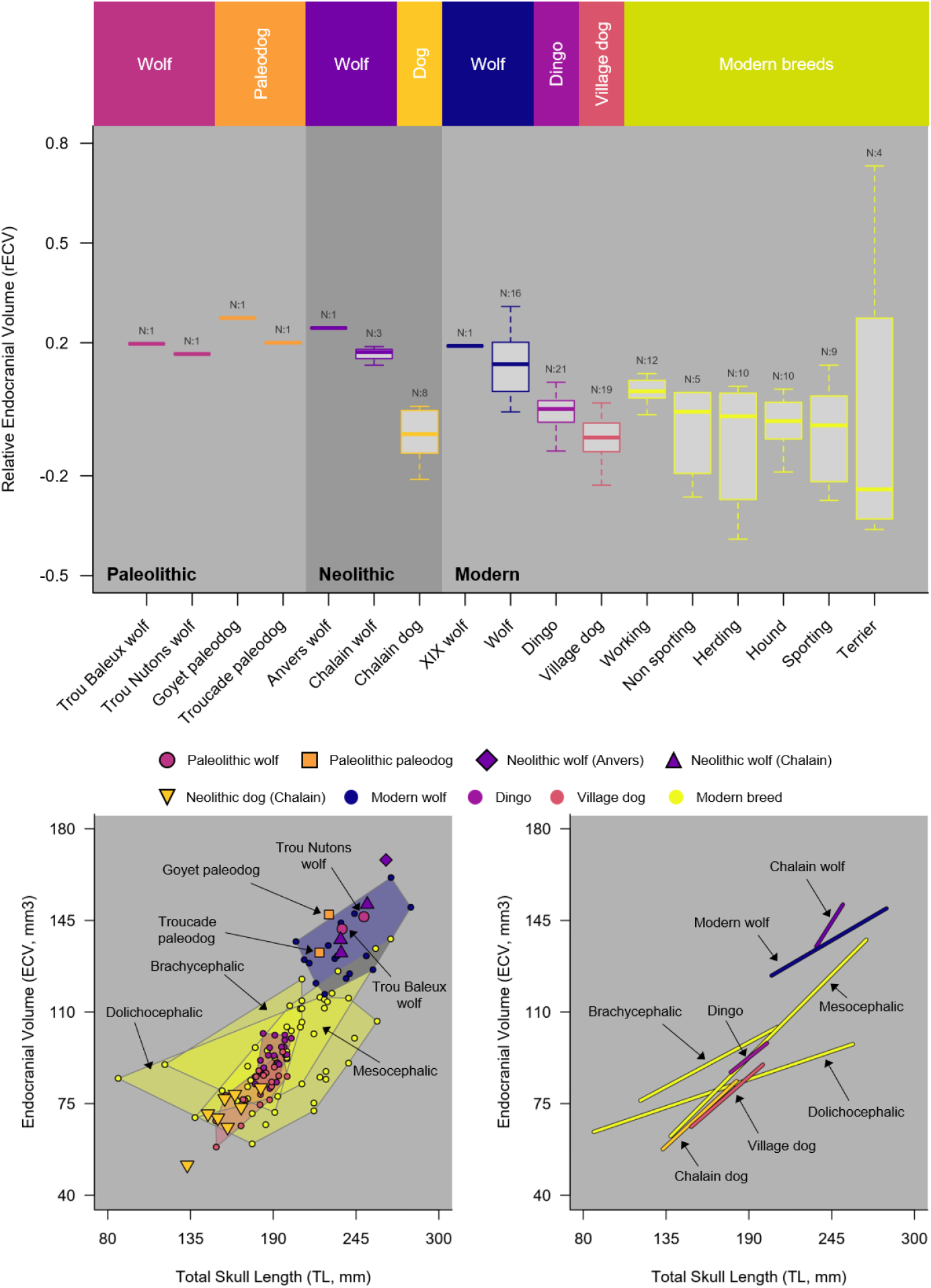
Variation in rECV (residuals from the regression of ECV on cranium length) across ancient (Palaeolithic and Neolithic) and modern wolves and dogs. (a) Boxplot of rECV values with modern dog breeds grouped by AKC functional categories. (b) Scatterplot of ECV against cranium length (TL). (c) Same scatterplot with only the slopes of the linear regression performed on groups with more than 2 specimens.

To compare the rECV across ancient and modern dogs and wolves, we excluded Toy dogs as well as Pugs and French Bulldogs as these very small and extremely brachycephalic dogs are clearly outliers. We found no significant rECV differences across dingoes, village dogs, and the functional groups of dog breeds (SI Table 8). Nevertheless, working dogs show, on average, higher rECV than other functional groups (Figure 5a). The rECV values of dingoes are similar to those of mesocephalic breeds of dogs of the same skull size, but larger than those of village dogs of the same skull size (Figure 5a, b,c). Finally, the Neolithic Chalain dogs show smaller rECV than dingoes and mesocephalic dogs in a lesser extent but similar rECV values than village dogs (Figure 5a,b,c).

### 4. Neurocranial anatomy in ancient and modern dogs and wolves

We compared neurocranial variation across ancient and modern wolves and dogs, using a scatterplot of their ECV against their cephalic index (CI) values (Figure 6), excluding Toy dogs, Pugs, and the French Bulldog. In this dataset, the CI showed no significant relationship with endocranial volume (F-statistic = 0.515, *Df* = 114, *P* = *0*.*474*). The scatterplot shows that Pleistocene wolf from Les Nutons and the protodogs from Goyet and La Baume Traucade cluster with modern and Neolithic wolves. Dingoes, village dogs, and the Chalain dogs cluster together, with the Chalain dogs showing greater similarity to village dogs and some mesocephalic herding breeds (e.g., Rough Collie, Australian Shepherd Dog) than to dingoes.

**Figure 6:**
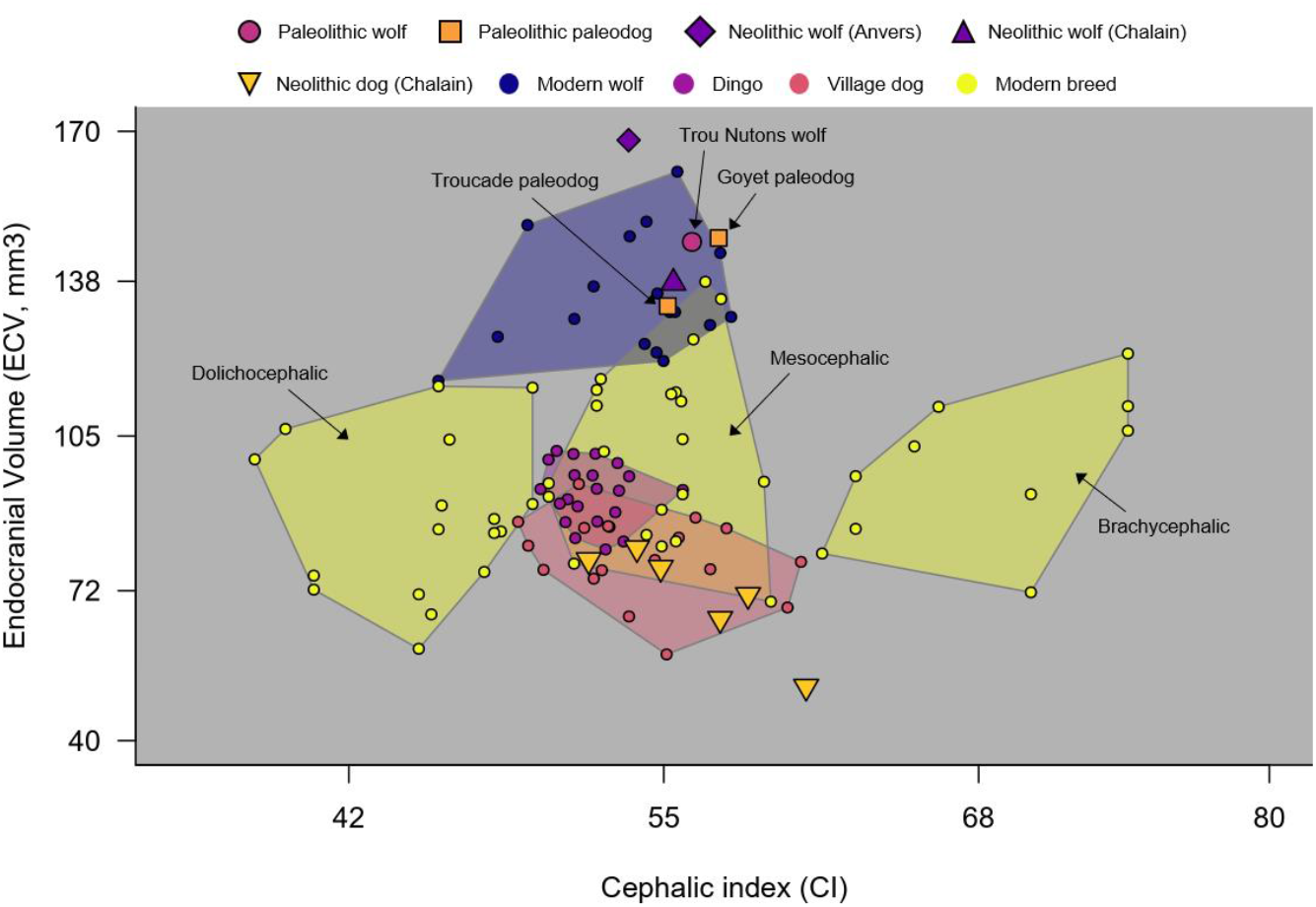
Neurocranial variation in modern and ancient wolves and dogs (excluding Toy dogs such as Pugs and French Bulldogs). Scatterplot of the Endocranial volume (ECV in mm^3^) against the cephalic index (CI).

## Discussion

Thanks to advances in endocast imaging (e.g., Czeibert et al., 2024), we were able to accurately estimate brain size in 22 archaeological and palaeontological canid specimens. This allowed us to track how brain size changed during different phases of the evolutionary history of dogs. We found that so-called “protodogs” from the Pleniglacial period in Belgium (Goyet) and the Late Glacial period in France (Baume Traucade), show no brain size reduction and possess a neurocranial anatomy most similar to Pleistocene, Neolithic and modern wolves. This would strongly support their identification as wolves, following previous studies which proposed that Western European Pleniglacial dogs are part of an ecophenotypic diversity of Pleistocene wolves that is now extinct (Boudadi-Maligne and Escarguel, 2014; Janssens et al., 2019), although this interpretation depends on the comparative framework and methods used (see Galeta et al., 2021). Yet, we found that the Goyet “protodog” shows a larger relative brain volume than Pleistocene, Neolithic, and modern wolves of comparable skull size. The behavioural flexibility to fit into the human social environment (Hare et al., 2002) or to access food resources, as suggested for captive wolves (Siciliano-Martina et al., 2022), could lead to an increase in brain size in these animals (Sol et al., 2008), as observed for small mammals adapting to urbanization (DePasquale et al., 2020). Recent neuroanatomic analysis of grey foxes of the fox-farm experiment (Trut, 1999), showed that grey foxes selected for tameness displayed a greater amount of grey matter in the frontal cortex (Hecht et al., 2021). Therefore, a brain size increases rather than a decrease, might be the response to expect during early process of intensification in human and canid interactions.

Our study confirms the 30% brain size reduction between modern wolves and dogs previously identified (Balcarcel et al., 2024; Garamszegi et al., 2023), with a threefold reduction in endocranial volume between the smallest- and the largest-brained breed of dogs. Despite this reduction we found that working dogs show greater ECV than dingoes, village dogs, and dog breeds, even when accounting for body size, suggesting that working dogs have larger brains than other functional groups. This trend differs from a recent study which reported working dogs as having the smallest rECV of all dogs (Balcarcel et al., 2024). This is probably due to the fact that we have more large dogs in the working group and a different statistical approach to compute the relative ECV. Nonetheless, our results tend to be in agreement with evidence that greater brain volume predicts cognitive performance and self-control (MacLean et al., 2014) and underlies the greater trainability observed in working dogs (Horschler et al., 2019; McGreevy et al., 2013).

We then found that brain size reduction between wolves and dogs is already established by 5,000-4,500 BP and is even more important than today since the mesocephalic dogs of Late Neolithic Chalain have a 46% smaller brain size than modern and Neolithic wolves, comparable to the brain size of small terrier and Toy breeds such as Pugs, Chihuahua, and Pekingese. This strongly suggests that selection for small dogs have begun in Europe during the Neolithic period in Europe, maybe before or at the same time as giant flock-guardian dogs, often regarded as the first morphotype to appear independently in different regions of the world (Parker et al., 2017). The dogs of Chalain exhibit a mesocephalic cranial morphology typical of free-ranging village dogs and a skull size close to terrier dogs like medium sized German Spitz, resembling other Neolithic dogs found in the UK and Switzerland. This pattern is consistent with the occurrence of a relatively homogeneous morphotype of small dogs in the Middle and Late Neolithic of Western Europe (Brassard et al., 2022; Harcourt, 1974; Horard-Herbin et al., 2014).

This drastic brain size reduction in these small Late Neolithic dogs suggests also accompanying behavioural changes. Indeed, recent studies have shown that small dogs tend to be more fearful, aggressive towards strangers, prone to barking (Ayrosa et al., 2022; McGreevy et al., 2013; Stone et al., 2016; Turcsán and Kubinyi, 2025) and less trainable (Serpell, 2017; Zapata et al., 2016). The latest neuroanatomical studies explain these behavioural differences by the reorganisation of the brain tissues induced by the change in brain size (Barton et al., 2025; Hecht et al., 2021b). Brain size decrease induces the reduction of the cortical brain along with an increase in subcortical tissues, whereas increases in brain size are associated with greater cortical and less subcortical tissues. This brain tissue reorganisation induced by size reduction means less cognitive abilities and more anxiety driven behaviours, while brain size increase means greater trainability and less social anxiety. This would suggest that the small brained Neolithic dogs of Chalain were likely more fearful, anxious, prone to barking and not very trainable, raising questions about their role in this Late Neolithic farming socio-ecosystem.

Zooarchaeologists have already proposed that Middle and Late Neolithic dogs in this region served polyvalent purposes, including symbolic and ornamental roles with no clear economic or utilitarian role (Arbogast et al., 2005). Their use in ritual feasting has also been suggested for Neolithic contexts in Italy (De Grozzi Mazzorin and Tagliacozzo, 1997). Given the strong neurocranial similarity between the Chalain dogs and present-day pariah-type village dogs, the latter may provide an analogy for their purpose, while their small brain size provides insight into their temperament. The dogs of Chalain were likely part of the Neolithic settlement socio-ecosystem like village dogs, scavenging food refuse and potentially used as a convenient source of meat, like in many parts of the world today with clear examples from Eastern and South Eastern Asia (e.g., (Avieli, 2011; Li et al., 2017; Podberscek, 2009). The dog bones from Chalain were found disconnected and among the other faunal remains consumed by the village settlers (Pétrequin, 1997), which would support this interpretation. What their small brain implies in term of anxious temperament and higher reactivity to novelty, lead us to consider that these small Neolithic village dogs could have served the purpose of alerting for any unexpected changes in the settlement surroundings. Like modern village dogs, their poor diet and harsh living conditions, may have also imposed metabolic constraints on their growth (Dickerson et al., 1997; Faust et al., 2021), contributing further to their small brain size. One might then ask whether this highly anxious and reactive temperament could have been one of the main targets of dog selection during the Neolithic.

Finally, we found that dingoes display an endocranial volume that is intermediate between the largest- and smallest-brained dogs, but greater than village dogs of the same skull size. This suggesting the reversibility of brain size reduction by domestication through feralization (Kruska, 2005), which has been observed in other species, such as mink (Pohle et al., 2023). This is consistent with previous large-scale studies comparing the relative brain size of dingoes with dogs and other wild canids, which suggest that brain size increases in dingoes could be linked to their evolution as apex predators in the Australian environment (Smith et al., 2018). Reduced metabolic constraints of placental predators in an environment of marsupial mammals along with the behavioural flexibility required by the cognitive challenges of the Australian environment, could have been strong evolutionary constraints as well (Hecht et al., 2023).

## Conclusion

Our results provide new evidence for changes in brain size during the evolutionary history of dogs. We found no evidence of brain size reduction in “protodogs” of the Upper Pleistocene. Instead, we raise the possibility that the intensification in humans and canids interaction may be due to a slight increase in brain size, reflecting the adaptation to the cognitive challenges of living in proximity to humans, or easier access to food resources. This study found however a dramatic 46% brain size reduction by 5,000-4,500 BP in Late Neolithic dogs compared to contemporaneous wolves, with brain volume close to that of recent Toy breeds, providing clear evidence for very early behavioural selection. Relying on the latest understanding in the link between neuroanatomy and dogs’ temperament, this drastic brain size reduction in Neolithic provides important clues for their potential use for alerting the settlement against threats, among other functions such as scavenging, convenient source of meat or hunting. Both interpretations require further testing with more samples of Palaeolithic, Mesolithic and Neolithic wolves and dogs from Europe. Furthermore, endocranial volume is only a proxy for brain anatomy that does not account for changes in brain proportions or more localised changes driven by natural and artificial selection for behavioural specialisation to specific anthropogenic environments (Hecht et al., 2019). Addressing this hypothesis, however, will require future studies to explored more aspects of complex anatomical features of the endocast and its integration with the skull (Schwab et al., 2023), using the latest developments in the quantitative methods for anatomy (Bardua et al., 2019). Such approach will allow the exploration and disentanglement of smaller scale anatomical changes associated with behavioural changes during different stages of dog domestication, including the initial transition from wild to domestic and later adaptation to the multitude of anthropogenic environments to which dogs have been exposed to over their long co-evolution with people.

## Acknowledgments

The authors warmly thank Arnaud Larralle (president of the FRGP Nouvelle-Aquitaine (Regional Federation of Private Guards of Nouvelle-Aquitaine) and Raymond Triquet (former Chairman of the standards commission of the FCI) for their help in retrieving wolf heads from the OFB. We would like to express our thanks to the Lons-Le-Saunier Museum (France) and Eve Neyret, curator of the conservation and research center (CCE) of Lons-Le Saunier, for their agreement and help to access the archaeological specimens of Chalain. We also would like to thanks the Municipality of Issirac (France), owner of the La Baume Traucade site, in the person of its Mayor Mr José Rieu, as well as Romain Franquet, Curator of the National Nature Reserve of the Gorges de l’Ardèche, and its Scientific Committee, for their authorisations. The support of the DRAC Occitanie was essential to this survey operation. Our sincere thanks go to Cyril Montoya (Deputy Regional Archaeology Curator), Denis Guilbeau (Head of Recent Archaeology) and Philippe Galant (Head of Ornate Caves and Underground Heritage).) as well as La Cité de la Préhistoire in Orgnac l’Aven: Patricia Guillermin (Heritage Curator, Director of La Cité de la Préhistoire) and Laurine Viel, Collections and Documentation Manager. Sandy Ingleby (Australian Museum) for access to dingo specimens. Richard Sabin and Phaedra Kokkini (NHM London) for access to historical New Guinea singing dog specimens. Brett Clark (NHM London) for assistance with micro-CT scanning at the NHM Imaging and Analysis Centre

