## Supplementaries for "Brain size reduction in dogs was already established at least by the Late Neolithic of western Europe, 5,000 years ago"

| Taxon | Period | AKC | Breeds | Phylogenetic clade | N | Collection curators |
|---|---|---|---|---|---|---|
| *Canis lupus lupus* | *21st century* France | NA | NA | NA | 58 | ONIRIS |
| *Canis lupus lupus* | *19th century* Belgium | NA | NA | NA | 1 | RBINS |
| *Canis lupus dingo* | *19th-20$^{th}$ Australia* | NA | NA | NA | 21 | Australian Museum Sydney |
| *Canis familiaris* Village dogs | *19th-20th* worldwide | | | | 19 | NHM London |
| *Canis familiaris* | *Modern (20th)* | Herding | Australian Sheperd, Border collie, Briard | UK rural, continental herder, New world | 11 | ELTE (Hungary) |
| *Canis familiaris* | *Modern (20th)* | Hound | Afghan hound, Basset Hound, Beagle, Borzoi, Greyhound, Rhodesian Ridgeback, Saluki | Mediterranean, Scent Hound, Sight Hound, European Mastiff | 10 | ELTE (Hungary) |
| *Canis familiaris* | *Modern (20th)* | Non-sporting | American Bulldog, Bulldog, Chow-chow, Dalmatian, French bulldog, Medium German Spitz | European Mastiff, Pinscher, Asian Spitz, Pointer Setter | 7 | ELTE (Hungary) |
| *Canis familiaris* | *Modern (20th)* | Sporting | English Cocker Spaniel, English Pointer, Golden retriever, Labrador retriever, Weimaraner | Spaniel, Pointer Setter, Retriever | 9 | ELTE (Hungary) |
| *Canis familiaris* | *Modern (20th)* | Terrier | Cairn Terrier, Fox Terrier, Staffordshire Bull Terrier, Welsh Terrier | Terrier | 6 | ELTE (Hungary) |
| *Canis familiaris* | *Modern (20th)* | Toy | Chihuahua, Pekingese, Pug | American Toy, Asian Toy, Toy Spitz | 6 | ELTE (Hungary) |
| *Canis familiaris* | *Modern (20th)* | Working | Doberman Pinscher, Greater Swiss Mountain dog, Siberian Husky, Cane Corso, Great Dane, Leon Berger, English Mastiff, Saint Bernard | European Mastiff, Pinscher, Alpine, Asian Spitz | 12 | ELTE (Hungary) |

SI Table 1: Modern dogs and wolves' samples used as comparative material for this study. AKC and Phylogenetic clades have been obtained from Garamzegui et al. (2023).

**SI Table 2:** Sample details for the archaeological dogs and wolves' studied

| Site | Country | Context | Specimen | Type | Taxon |
|---|---|---|---|---|---|
| Chalain 4 (F et G) fouilles Pétrequin | France | Lake dwelling site | DF65D | cranium | Canis lupus |
| Chalain 4 (F et G) | France | Lake dwelling site | HF60B | cranium | Canis lupus |
| Chalain 4 (F et G) | France | Lake dwelling site | EF62D | cranium | Canis lupus |
| Chalain 4 (F et G) | France | Lake dwelling site | DF64C | braincase | Canis lupus |
| Chalain 4 (F et G) | France | Lake dwelling site | BF65F | cranium | Canis lupus |
| Chalain "fonds ancien" | France | Lake dwelling site | M0346_00726 | cranium | Canis lupus |
| Chalain "fonds ancien" | France | Lake dwelling site | M0346_00728 | cranium | Canis lupus |
| Chalain "fonds ancien" | France | Lake dwelling site | M0346_00621 | cranium | Canis lupus |
| Chalain "fonds ancien" | France | Lake dwelling site | M0346_01040 | cranium | Canis lupus |
| Chalain "fonds ancien" | France | Lake dwelling site | M0346_00725 | cranium | Canis lupus |
| Chalain "fonds ancien" | France | Lake dwelling site | M0346_00730 | cranium | Canis lupus |
| Chalain "fonds ancien" | France | Lake dwelling site | M0346_00732 | braincase | Canis lupus |
| Chalain "fonds ancien" | France | Lake dwelling site | M0346_00623 | cranium | Canis lupus |
| Goyet | Belgium | Cave deposit | DP2860 | cranium | *Canis lupus sp.* |
| Trou des Nutons | Belgium | Cave deposit | DP2559 | cranium | *Canis lupus sp.* |
| Grands Malades | Belgium | Cave deposit | | cranium | *Canis lupus sp.* |
| La Baume Traucade | France | Cave deposit | | cranium | *Canis lupus sp.* |
| Trou Balleux | Belgium | Cave deposit | ULg | cranium | *Canis lupus sp.* |
| Antwerpen | Belgium | open-air site | | cranium | *Canis lupus* |

| Status | Chronology | Culture | Curation | References |
|---|---|---|---|---|
| wolf | 3010-2995 BC | Late Neolithic, Early Clairvaux | CCE Lons Le Saunier | Pétrequin et al. 1997 |
| dog | 3010-2995 BC | idem | idem | idem |
| wolf | 3010-2995 BC | idem | idem | idem |
| wolf | 3010-2995 BC | idem | idem | idem |
| wolf | 3010-2995 BC | idem | idem | idem |
| dog | circa 3000-2650 BC | Late Neolithic, Early Clairvaux to Chalain/ Auvernier Cordée | idem | idem |
| dog | circa 3000-2650 BC | idem | idem | idem |
| dog | circa 3000-2650 BC | idem | idem | idem |
| dog | circa 3000-2650 BC | idem | idem | idem |
| dog | circa 3000-2650 BC | idem | idem | idem |
| dog | circa 3000-2650 BC | idem | idem | idem |
| dog | circa 3000-2650 BC | idem | idem | idem |
| wolf | circa 3000-2650 BC | idem | idem | idem |
| Protodog | 35,500 cal BP | Aurignacian | RBINS | Germonpré et al. 2009, 2012 |
| Pleistocene wolf | 26,000 cal BP | Gravettian | RBINS | Germonpré et al. 2009 |
| Pleistocene wolf | Pleniglacial | Mousterian? | RBINS | Germonpré et al. 2009 |
| Protodog | 15,500 cal BP | ? | Musée d'Orgnac | Germonpré et al. 2025 |
| Holocene wolf | Postglacial | ? | RBINS | Germonpré et al. 2009 |
| Holocene wolf | Neolithic | ? | RBINS | Hasse, 1909 |

| Taxon | N | Collection curators | CT parameters |
|---|---|---|---|
| *Canis lupus lupus* | *60* | ONIRIS (France) | ONIRIS: SOMATOM Go.Up CT scanner (Siemens Healthineers GmbH, Germany). Helical acquisition was acquired with the following parameters: 130 kVp, modulation of the mAs via the Care Dose 4D function, slice thickness of 0.6 mm with 0.4 mm increment, tube revolution time of 1.5s and spiral pitch factor of 0.35. Bone and soft tissue reconstruction algorithms were applied |
| *Canis lupus lupus* | *1* | RBINS ULg (Belgium) | Universitair Ziekenhuis Leuven campus, Helical CT scan (Siemens Somatom Sensation 64) with a voxel size of 1 mm isotropic, matrix 512 x 512. |
| *Canis sp* | *5* | | |
| *Canis lupus dingo* | *21* | Australian Museum Sydney | kV 195 uA 44 Projections/Frames 2200/2 Exposure (Milliseconds) 354 Gain (db) 24 |
| *Canis familiaris* Village dogs | *19* | NHM London (UK) | Nikon Metrology HMX ST 225 at the Imaging and Analysis Centre at the Natural History Museum, London, UK, with scattering artefacts reduced using a 0.1 mm copper filter |
| *Canis familiaris* | *64* | ELTE (Hungary) | Siemens Somatom Definition AS+ CT machine (Siemens, Erlangen, Germany; 170 mAs, 140 kV, pixel size 0.323 × 0.322mm, slice thickness 0.6mm, with a v80u bone kernel). |
| *Canis sp. (Baume Traucade)* | *1* | Musée d'Orgnac (France) | PIXANIM INRAE imaging plateform Siemens Somatom Definition AS+ CT machine (Siemens, Erlangen, Germany; 170 mAs, 140 kV, pixel size 0.323 × 0.322mm, slice thickness 0.6mm, with a v80u bone kernel). |
| *Canis lupus lupus* | *15* | CCE Lons-Le-Saunier (France) | |
| *Canis familiaris* | | | |

**SI Table 3:** Details for the different CT parameters used across the different wolves and dogs' collection studied.

| Specimen | Status | Period | TL | Reference |
|---|---|---|---|---|
| Maldidier | « Pleistowolf » | 30,200 cal BP | 254 | (Boudadi-Maligne, 2010) |
| Anabar | « Pleistowolf » | Pleistocene | 268 | (Germonpré et al., 2009) |
| Avdeevo_911 | Large *Canis* | Pleistocene | 246 | (Germonpré et al., 2009) |
| Goyet | « Protodog » | 36,100 cal BP | 227 | (Germonpré et al., 2009) |
| Mezin_5488 | « Pleistowolf » | Pleistocene, post-LGM | 256 | (Germonpré et al., 2009) |
| Mezin_5469 | « Pleistowolf » | Pleistocene, Post-LGM | 254 | (Germonpré et al., 2009) |
| Trou des Nutons | « Pleistowolf » | 26,000 cal BP | 261 | (Germonpré et al., 2009) |
| Kostenki-17 | « Pleistowolf » | 39,500 cal BP | 265 | (Germonpré et al., 2012; Germonpré & Sablin, 2017)) |
| Předmostí _1_CR | « Pleistowolf » | 28,500 cal BP | 277 | (Germonpré et al., 2012, 2017b) |
| Předmostí _1924 | Large *Canis* | 28,500 cal BP | 274 | (Germonpré et al., 2012, 2017b) |
| Předmostí _1062 | Large *Canis* | 28,500 cal BP | 264 | (Germonpré et al., 2012, 2017b) |
| Předmostí _1060 | « Protodog » | 28,500 cal BP | 238 | (Germonpré et al., 2012; 2017b) |
| Předmostí _1069 | « Protodog » | 28,500 cal BP | 236 | (Germonpré et al., 2012; 2017b) |
| Předmostí -CR | « Protodog » | 28,500 cal BP | 232 | (Germonpré et al., 2012; 2017b) |
| Předmostí _1061 | « Protodog » | 28,500 cal BP | 228 | (Germonpré et al., 2012; 2017b) |
| Badyarikha | Large *Canis* | 29,800 cal BP | 258 | (Germonpré et al., 2017a) |
| Tirekhtyakh | « Pleistowolf » | >50,000 cal BP | 254 | (Germonpré et al., 2017a) |
| Ulakhan_Sular | « Protodog » | 16,900 cal BP | 206 | (Germonpré et al., 2017a) |
| Baume Traucade | « Protodog » | 15,500 cal BP | 221 | (Germonpré et al., 2025) |
| Mezherichi_4493 | « Protodog » | 17,900 cal BP | 226 | (Germonpré et al., 2012; Chu et al., 2025) |
| Razboinichya | « Protodog » | 33,400 cal BP | 211 | (Ovodov et al., 2011) |
| Mezin_5490 | « Protodog » | Pleistocene, post-LGM | 226 | (Pidoplichko and Allsworth-Jones, 1998) |
| Eliseevichi | « Protodog » | 16,900 cal BP | 240 | (Sablin and Khlopachev, 2002; Sablin, 2008) |
| Eliseevichi | « Protodog » | 16,000 cal BP | 256 | (Sablin and Khlopachev, 2002; Sablin, 2008) |
| Chalain_621 | dog | Neolithic 5,000-4,500 cal BP | 169 | This_study |
| Chalain_725 | dog | Neolithic 5,000-4,500 cal BP | 153 | This_study |
| Chalain_726 | dog | Neolithic 5,000-4,500 cal BP | 158 | This_study |
| Chalain _728 | dog | Neolithic | 165 | This_study |

| | | | | |
|---|---|---|---|---|
| | | 5,000-4,500 cal BP | | |
| Chalain _1040 | dog | Neolithic 5,000-4,500 cal BP | 147 | This_study |
| Chalain _C277 | dog | Neolithic 5,000-4,500 cal BP | 182 | This_study |
| Chalain _C2229 | dog | Neolithic 5,000-4,500 cal BP | 133 | This_study |
| Chalain _HF60B | dog | Neolithic 5,000-4,500 cal BP | 160 | This_study |
| Easton_Down | dog | Neolithic | 168 | (Harcourt, 1974) |
| Dowel_Cave | dog | Neolithic | 178 | (Harcourt, 1974) |
| Maiden_Castle_I | dog | Neolithic | 176 | (Harcourt, 1974) |
| Maiden_Castle_II | dog | Neolithic | 177 | (Harcourt, 1974) |
| Windmill_Hill_I | dog | Neolithic | 174 | (Harcourt, 1974) |
| Windmill_Hill_II | dog | Neolithic | 169 | (Harcourt, 1974) |
| NeoDog_Switz1 | dog | Neolithic | 164 | (Janssens et al., 2019) |
| NeoDog_Switz2 | dog | Neolithic | 173 | (Janssens et al., 2019) |
| NeoDog_Switz3 | dog | Neolithic | 159 | (Janssens et al., 2019) |
| NeoDog_Switz4 | dog | Neolithic | 155 | (Janssens et al., 2019) |
| NeoDog_Switz5 | dog | Neolithic | 151 | (Janssens et al., 2019) |
| NeoDog_Switz6 | dog | Neolithic | 173 | (Janssens et al., 2019) |
| NeoDog_Switz7 | dog | Neolithic | 168 | (Janssens et al., 2019) |
| NeoDog_Switz8 | dog | Neolithic | 157 | (Janssens et al., 2019) |
| NeoDog_Switz9 | dog | Neolithic | 157 | (Janssens et al., 2019) |
| NeoDog_Switz10 | dog | Neolithic | 163 | (Janssens et al., 2019) |
| NeoDog_Switz11 | dog | Neolithic | 175 | (Janssens et al., 2019) |
| NeoDog_Switz12 | dog | Neolithic | 156 | (Janssens et al., 2019) |
| NeoDog_Switz13 | dog | Neolithic | 170 | (Janssens et al., 2019) |
| NeoDog_Switz14 | dog | Neolithic | 172 | (Janssens et al., 2019) |
| NeoDog_Switz15 | dog | Neolithic | 160 | (Janssens et al., 2019) |
| NeoDog_Switz16 | dog | Neolithic | 173 | (Janssens et al., 2019) |
| NeoDog_Switz17 | dog | Neolithic | 166 | (Janssens et al., 2019) |
| NeoDog_Switz18 | dog | Neolithic | 165 | (Janssens et al., 2019) |
| NeoDog_Switz19 | dog | Neolithic | 146 | (Janssens et al., 2019) |
| NeoDog_Switz20 | dog | Neolithic | 162 | (Janssens et al., 2019) |
| NeoDog_Switz21 | dog | Neolithic | 166 | (Janssens et al., 2019) |
| NeoDog_Switz22 | dog | Neolithic | 162 | (Janssens et al., 2019) |
| NeoDog_Switz23 | dog | Neolithic | 156 | (Janssens et al., 2019) |
| NeoDog_Switz24 | dog | Neolithic | 158 | (Janssens et al., 2019) |
| NeoDog_Switz25 | dog | Neolithic | 166 | (Janssens et al., 2019) |
| NeoDog_Switz26 | dog | Neolithic | 153 | (Janssens et al., 2019) |
| NeoDog_Switz27 | dog | Neolithic | 159 | (Janssens et al., 2019) |
| NeoDog_Switz28 | dog | Neolithic | 155 | (Janssens et al., 2019) |
| NeoDog_Switz29 | dog | Neolithic | 158 | (Janssens et al., 2019) |
| NeoDog_Switz30 | dog | Neolithic | 171 | (Janssens et al., 2019) |
| NeoDog_Switz31 | dog | Neolithic | 157 | (Janssens et al., 2019) |
| NeoDog_Switz32 | dog | Neolithic | 157 | (Janssens et al., 2019) |

| | | | | |
|---|---|---|---|---|
| NeoDog_Switz33 | dog | Neolithic | 150 | (Janssens et al., 2019) |
| NeoDog_Switz34 | dog | Neolithic | 173 | (Janssens et al., 2019) |
| NeoDog_Switz35 | dog | Neolithic | 176 | (Janssens et al., 2019) |
| NeoDog_Switz36 | dog | Neolithic | 176 | (Janssens et al., 2019) |
| NeoDog_Switz37 | dog | Neolithic | 144 | (Janssens et al., 2019) |
| NeoDog_Switz38 | dog | Neolithic | 152 | (Janssens et al., 2019) |
| NeoDog_Switz39 | dog | Neolithic | 172 | (Janssens et al., 2019) |
| Muge_Dog | dog | Mesolithic 7,000 cal BP | 189 | (Detry and Cardoso, 2010) |
| Zhokhov_Dd140 | dog | Mesolithic, 9,000 cal BP | 196 | (Pitulko and Kasparov, 2017) |
| Zhokhov_Dd141 | dog | Mesolithic, 9,000 cal BP | 219 | (Pitulko and Kasparov, 2017) |
| Trou Balleux | Wolf | Postglacial | 238 | Germonpré et al. 2009 |

**SI Table 4:** Cranium length (TL) in mm of Pleistocene, Mesolithic and Neolithic *Canis* sp. collected from the literature (References).

| ECV (mm³) | N | Mean | sd | Min | Max |
|---|---|---|---|---|---|
| Pleistocene Wolves | 2 | 159.51 | 18.53 | 146.41 | 172.62 |
| Protodogs | 2 | 139.98 | 10.27 | 132.72 | 147.25 |
| Wolves (France) | 58 | 133.71 | 9.479 | 111.73 | 157.65 |
| Dingoes | 21 | 92.76 | 6.355 | 80.79 | 101.82 |
| Village_dogs | 18 | 79.94 | 8.793 | 58.40 | 94.75 |
| Herding | 11 | 92.54 | 14.911 | 72.22 | 117.16 |
| Hound | 10 | 94.94 | 11.443 | 76.85 | 113.93 |
| Non-sporting | 7 | 80.31 | 10.459 | 69.66 | 92.58 |
| Sporting | 9 | 98.31 | 15.719 | 76 | 114.83 |
| Terrier | 6 | 82.93 | 22.229 | 59.61 | 120.15 |
| Toy | 6 | 60.14 | 14.548 | 47.63 | 84.73 |
| Working | 12 | 111.18 | 17.251 | 85.09 | 137.91 |
| Chalain_dogs | 7 | 71.03 | 9.311 | 51.52 | 81.08 |
| Neolithic_wolves | 6 | 147.37 | 9.58 | 132.58 | 151.12 |

| TL (mm) | N | Mean | sd | Min | Max |
|---|---|---|---|---|---|
| Pleistocene Wolves | 1 | 250.21 | | | |
| Protodogs | 2 | 223.85 | 4.45 | 220.7 | 227 |
| Wolves (France) | 58 | 235.13569 | 19.58 | 205.1 | 281.38 |
| Dingoes | 21 | 190.45238 | 6.98 | 177.97 | 202.03 |
| Village_dogs | 18 | 179.36637 | 14.00 | 151.81 | 199.24 |
| Herding | 11 | 205.80000 | 27.97 | 156.00 | 240.00 |
| Hound | 10 | 204.60000 | 30.18 | 173.00 | 259.00 |
| Non-sporting | 7 | 174.00000 | 37.13 | 118.00 | 213.00 |
| Sporting | 9 | 211.11111 | 19.04 | 166.00 | 225.00 |
| Terrier | 6 | 156.00000 | 46.19 | 87.00 | 185.00 |
| Toy | 6 | 88.66667 | 11.64 | 76.00 | 102.00 |
| Working | 12 | 216.16667 | 27.14 | 178.00 | 268.00 |
| Chalain_dogs | 7 | 158.13500 | 14.68 | 132.87 | 181.77 |
| Neolithic_wolves | 5 | 248.89 | 13.52 | 234.73 | 264.89 |

**Supplementary Table 5**: Basic Statistics (Mean, Standard deviation (sd), maximum and minimum) for ECV ($mm^3$) and total skull length measurement (TL in mm) from modern and Chalain's wolves and dogs collected for this study.

**SI Figure 1:** Box plot of ECV values (mm$^3$) of ancient and modern wolves and dogs are played in a descending order and where all the dog's breeds are displayed. Chalain (CHA) wolves (w) and dogs (d) have there box plot background in light green.

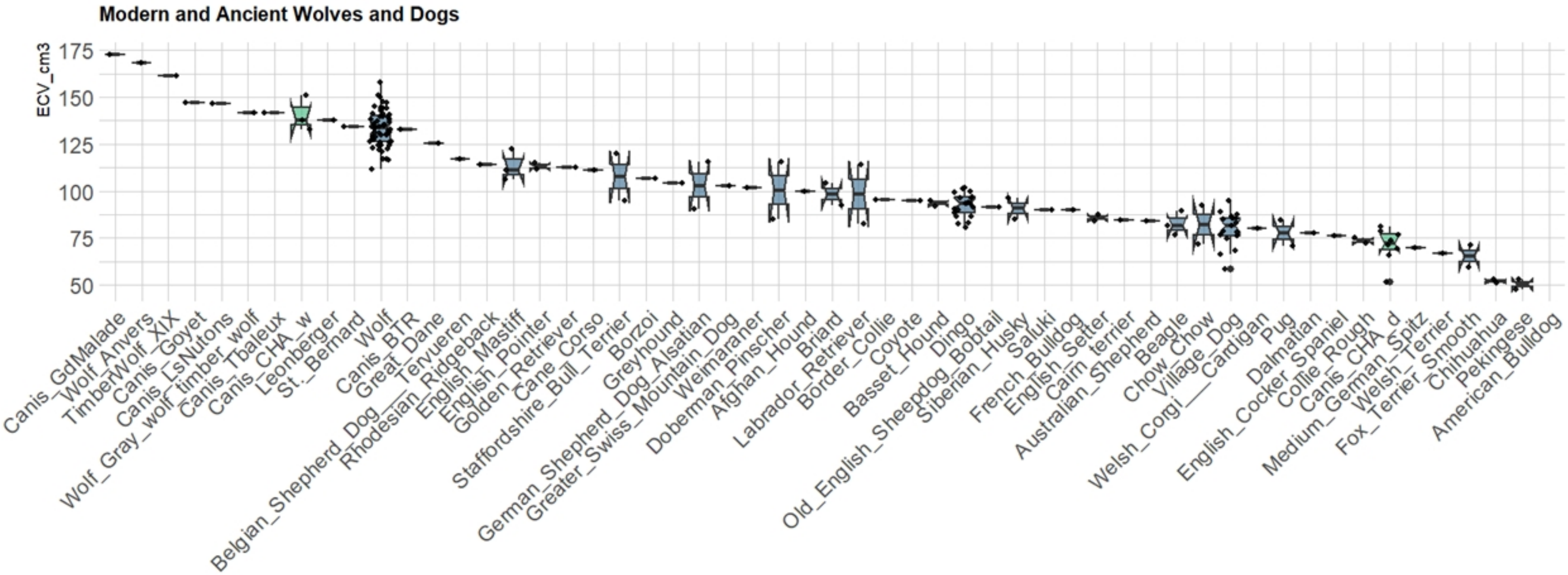

**SI Table 6:** Pairwise Comparison of cranium length (TL) between wolves and dogs' groups using Dunn's test with Bonferroni correction. Z values and adjusted p-values are reported. *: p values statistically significant. Dog_Bree: Dog breeds, Meso_dog: Mesolithic dogs, Mod_Wolf: Modern wolves, Neo_dog: Neolithic dogs, Neo_Wolf: Neolithic wolves, Plei_Wol: Pleistocene wolves.

| Col Mean- Row Mean | Dingo | Dog_Bree | Meso_dog | Mod_Wolf | Neo_dog | Neo_Wolf | Plei_Wol | Protodog |
|---|---|---|---|---|---|---|---|---|
| Dog_Bree | -0.046459 | | | | | | | |
| | 1.0000 | | | | | | | |
| Meso_dog | -0.462214 | -0.461594 | | | | | | |
| | 1.0000 | 1.0000 | | | | | | |
| Modern_W | -3.282099 | -3.831939 | -1.254371 | | | | | |
| | **0.0185*** | **0.0023*** | 1.0000 | | | | | |
| Neo_dog | 4.101467 | 5.604406 | 2.262727 | 7.635900 | | | | |
| | **0.0007*** | **0.0000*** | 0.4257 | **0.0000*** | | | | |
| Neo_Wolf | -2.604304 | -2.753072 | -1.383892 | -0.442526 | -5.030644 | | | |
| | 0.1657 | 0.1063 | 1.0000 | 1.0000 | **0.0000*** | | | |
| Plei_Wol | -4.089448 | -4.622005 | -1.850632 | -1.084933 | -7.937171 | -0.345536 | | |
| | **0.0008*** | **0.0001*** | 1.0000 | 1.0000 | **0.0000*** | 1.0000 | | |
| Protodog | -2.621851 | -2.949887 | -1.027879 | 0.323640 | -6.275919 | 0.652197 | 1.300885 | |
| | 0.1574 | 0.0572 | 1.0000 | 1.0000 | **0.0000*** | 1.0000 | 1.0000 | |
| Vill_dog | 1.362046 | 1.672726 | 1.153366 | 4.499217 | -2.342273 | 3.436347 | 5.182918 | 3.742627 |
| | 1.0000 | 1.0000 | 1.0000 | **0.0001*** | 0.3450 | **0.0106*** | **0.0000*** | **0.0033*** |

| Pairwise comparisons | Z | P.adjusted |
|---|---|---|
| Chalain_dog - Chalain_wolf | -3.91265636363207 | **0.00356016928406724*** |
| Chalain_dog - Dingo | -2.3330174868968 | 0.766241874164015 |
| Chalain_wolf - Dingo | 2.72121707326357 | 0.253663937688132 |
| Chalain_dog - Herding | -1.88068410664923 | 1 |
| Chalain_wolf - Herding | 2.72515846907974 | 0.250655461934549 |
| Dingo - Herding | 0.256399318327395 | 1 |
| Chalain_dog - Hound | -2.12343793130169 | 1 |
| Chalain_wolf - Hound | 2.49383903866111 | 0.492842357635963 |
| Dingo - Hound | -0.098710574578668 | 1 |
| Herding - Hound | -0.305210390657244 | 1 |
| Chalain_dog - Non_sporting | -0.654868007803616 | 1 |
| Chalain_wolf - Non_sporting | 3.11592335081808 | 0.0715142721083587 |
| Dingo - Non_sporting | 1.19767090705179 | 1 |
| Herding - Non_sporting | 0.928038655616061 | 1 |
| Hound - Non_sporting | 1.15734295887529 | 1 |
| Chalain_dog - Sporting | -2.16207340382069 | 1 |
| Chalain_wolf - Sporting | 2.39745306893592 | 0.643870333658675 |
| Dingo - Sporting | -0.203985065925776 | 1 |
| Herding - Sporting | -0.393132242305254 | 1 |
| Hound - Sporting | -0.094334658292413 | 1 |
| Non_sporting - Sporting | -1.21419682098676 | 1 |
| Chalain_dog - Terrier | -0.916999993006443 | 1 |
| Chalain_wolf - Terrier | 3.04571295250568 | 0.0905304392533499 |
| Dingo - Terrier | 1.02411395230349 | 1 |
| Herding - Terrier | 0.746065212367046 | 1 |
| Hound - Terrier | 0.991480738959439 | 1 |
| Non_sporting - Terrier | -0.201318479808516 | 1 |
| Sporting - Terrier | 1.05368780857331 | 1 |
| Chalain_dog - Toy | 0.261776066379617 | 1 |
| Chalain_wolf - Toy | 3.94601803909541 | **0.00309900356087902*** |
| Dingo - Toy | 2.3993526882539 | 0.64053938771394 |
| Herding - Toy | 2.00042526047685 | 1 |
| Hound - Toy | 2.22427425036403 | 1 |
| Non_sporting - Toy | 0.850011359191514 | 1 |
| Sporting - Toy | 2.26157383303539 | 0.92522610278636 |
| Terrier - Toy | 1.10264403748853 | 1 |
| Chalain_dog - Village_Dog | -0.837592601022034 | 1 |
| Chalain_wolf - Village_Dog | 3.69549516194767 | **0.0085588877237668*** |
| Dingo - Village_Dog | 1.94645647535657 | 1 |
| Herding - Village_Dog | 1.37479215735901 | 1 |
| Hound - Village_Dog | 1.67456411532535 | 1 |
| Non_sporting - Village_Dog | 0.0404226041727222 | 1 |
| Sporting - Village_Dog | 1.72386446203679 | 1 |
| Terrier - Village_Dog | 0.303702640946119 | 1 |
| Toy - Village_Dog | -1.05572822805079 | 1 |

| | | |
|---|---|---|
| Chalain_dog - Wolf | -6.34149732306748 | **8.8741657755398e-09*** |
| Chalain_wolf - Wolf | 0.438710703004136 | 1 |
| Dingo - Wolf | -5.58798459397877 | **8.95908069684108e-07*** |
| Herding - Wolf | -4.61409958108435 | **0.000153973214531321*** |
| Hound - Wolf | -4.0411815311241 | **0.00207412018741481*** |
| Non_sporting - Wolf | -4.32802573613703 | **0.000586762380770414*** |
| Sporting - Wolf | -3.74076720478626 | **0.00715491614179657*** |
| Terrier - Wolf | -4.42001494387661 | **0.000384906896704768*** |
| Toy - Wolf | -5.90567572440629 | **1.36969638116099e-07*** |
| Village_Dog - Wolf | -7.71932772679872 | **4.56084959899348e-13*** |
| Chalain_dog - Wolf_Antwerpen | -2.89965281877703 | 0.145694713985621 |
| Chalain_wolf - Wolf_Antwerpen | -0.369503055115838 | 1 |
| Dingo - Wolf_Antwerpen | -2.05781130322116 | 1 |
| Herding - Wolf_Antwerpen | -2.10793588356338 | 1 |
| Hound - Wolf_Antwerpen | -1.97205735872464 | 1 |
| Non_sporting - Wolf_Antwerpen | -2.46677265273098 | 0.531713867361561 |
| Sporting - Wolf_Antwerpen | -1.92105277294767 | 1 |
| Terrier - Wolf_Antwerpen | -2.38890254588345 | 0.659052586083969 |
| Toy - Wolf_Antwerpen | -2.97829057557648 | 0.113045805819918 |
| Village_Dog - Wolf_Antwerpen | -2.65359541517749 | 0.310593079897694 |
| Wolf - Wolf_Antwerpen | -0.680571525776557 | 1 |
| Chalain_dog - Working | -3.43307983474064 | **0.0232738877174976*** |
| Chalain_wolf - Working | 1.67607458252667 | 1 |
| Dingo - Working | -1.6515999723448 | 1 |
| Herding - Working | -1.66042464378091 | 1 |
| Hound - Working | -1.30728113166956 | 1 |
| Non_sporting - Working | -2.24247258621936 | 0.972302960309866 |
| Sporting - Working | -1.17108638202644 | 1 |
| Terrier - Working | -2.14348558281983 | 1 |
| Toy - Working | -3.41670924654851 | **0.0247193441168995*** |
| Village_Dog - Working | -3.29224838533487 | **0.0387620177425112*** |
| Wolf - Working | 2.59652437385102 | 0.367271673724568 |
| Wolf_Antwerpen - Working | 1.44938416433422 | 1 |

SI Table 7: **Pairwise Comparison of ECV ($mm^3$) between Neolithic and modern wolves and dogs' groups** using Dunn's test with Bonferroni correction. Z values and adjusted p-values are reported. *: p values statistically significant.

| Pairwise comparisons | Z | P.adjusted |
|---|---|---|
| Chalain_dog - Chalain_wolf | -3.90681006569416 | **0.00715447964876866*** |
| Chalain_dog - Dingo | -2.23873899450228 | 1 |
| Chalain_wolf - Dingo | 2.77826759365225 | 0.418069377284776 |
| Chalain_dog - Herding | -1.80831146580021 | 1 |
| Chalain_wolf - Herding | 2.77071189814924 | 0.427894280384918 |
| Dingo - Herding | 0.241510234087133 | 1 |
| Chalain_dog - Hound | -2.04440892268089 | 1 |
| Chalain_wolf - Hound | 2.54477292374913 | 0.836518486808549 |
| Dingo - Hound | -0.103092417759201 | 1 |
| Herding - Hound | -0.296380480767292 | 1 |
| Chalain_dog - Non_sporting | -0.400486999049827 | 1 |
| Chalain_wolf - Non_sporting | 3.14191303551654 | 0.128403629438156 |
| Dingo - Non_sporting | 1.25542728104742 | 1 |
| Herding - Non_sporting | 1.01906050905172 | 1 |
| Hound - Non_sporting | 1.22463037544462 | 1 |
| Chalain_dog - Sporting | -2.09713577871563 | 1 |
| Chalain_wolf - Sporting | 2.43884714633349 | 1 |
| Dingo - Sporting | -0.223101812281292 | 1 |
| Herding - Sporting | -0.397748093743828 | 1 |
| Hound - Sporting | -0.107246836050848 | 1 |
| Non_sporting - Sporting | -1.28764379485675 | 1 |
| Chalain_dog - Terrier | -0.886144374365813 | 1 |
| Chalain_wolf - Terrier | 3.06368191638581 | 0.167252893994318 |
| Dingo - Terrier | 0.975494637498783 | 1 |
| Herding - Terrier | 0.712638406657979 | 1 |
| Hound - Terrier | 0.951157537598069 | 1 |
| Non_sporting - Terrier | -0.36146624868498 | 1 |
| Sporting - Terrier | 1.02543587691897 | 1 |
| Chalain_dog - Village_Dog | -0.796914204666674 | 1 |
| Chalain_wolf - Village_Dog | 3.71672055192721 | **0.0154396478433988*** |
| Dingo - Village_Dog | 1.87689175996044 | 1 |
| Herding - Village_Dog | 1.3312829728876 | 1 |
| Hound - Village_Dog | 1.62249531597202 | 1 |
| Non_sporting - Village_Dog | -0.164733526935917 | 1 |
| Sporting - Village_Dog | 1.68825466826308 | 1 |
| Terrier - Village_Dog | 0.304728762328906 | 1 |
| Chalain_dog - Wolf | -6.28616615823636 | **2.48932275320644e-08*** |
| Chalain_wolf - Wolf | 0.467246277242205 | 1 |
| Dingo - Wolf | -5.66009582453312 | **1.15735698684214e-06*** |
| Herding - Wolf | -4.65301963441751 | **0.00025023845151715*** |
| Hound - Wolf | -4.08983989192817 | **0.00330228364286432*** |
| Non_sporting - Wolf | -4.10926227252511 | **0.00303647687879598*** |
| Sporting - Wolf | -3.77068816234784 | **0.0124540469858356*** |
| Terrier - Wolf | -4.41025375869925 | **0.000789859093147023*** |
| Village_Dog - Wolf | -7.70529269492309 | **9.98659563613369e-13*** |

| | | |
|---|---|---|
| Chalain_dog - Wolf_Antwerpen | -2.9625226078259 | 0.233423991759792 |
| Chalain_wolf - Wolf_Antwerpen | -0.430680339891216 | 1 |
| Dingo - Wolf_Antwerpen | -2.16123134677187 | 1 |
| Goyet_Protodog - Wolf_Antwerpen | -0.101159308559909 | 1 |
| Herding - Wolf_Antwerpen | -2.20397747072273 | 1 |
| Hound - Wolf_Antwerpen | -2.07137977739426 | 1 |
| Non_sporting - Wolf_Antwerpen | -2.59113995477434 | 0.731788136689562 |
| Sporting - Wolf_Antwerpen | -2.01424904394798 | 1 |
| Terrier - Wolf_Antwerpen | -2.46606726246999 | 1 |
| Village_Dog - Wolf_Antwerpen | -2.73530078943896 | 0.476773223548555 |
| Wolf - Wolf_Antwerpen | -0.767369171584203 | 1 |
| Chalain_dog - Wolf_XIX | -2.9432541681002 | 0.248457963097621 |
| Chalain_wolf - Wolf_XIX | -0.412981147840892 | 1 |
| Dingo - Wolf_XIX | -2.14126396521348 | 1 |
| Herding - Wolf_XIX | -2.1844102776816 | 1 |
| Hound - Wolf_XIX | -2.05189360827484 | 1 |
| Non_sporting - Wolf_XIX | -2.57286030782708 | 0.771593600916214 |
| Sporting - Wolf_XIX | -1.99486055047683 | 1 |
| Terrier - Wolf_XIX | -2.44714603027712 | 1 |
| Village_Dog - Wolf_XIX | -2.71538100599728 | 0.506426645404743 |
| Wolf - Wolf_XIX | -0.747102931152928 | 1 |
| Wolf_Antwerpen - Wolf_XIX | 0.0144513297942727 | 1 |
| Chalain_dog - Working | -3.35072294280965 | 0.0616596883469116 |
| Chalain_wolf - Working | 1.7281780502822 | 1 |
| Dingo - Working | -1.65596531584723 | 1 |
| Herding - Working | -1.65093327408241 | 1 |
| Hound - Working | -1.30703857640279 | 1 |
| Non_sporting - Working | -2.22419846765737 | 1 |
| Sporting - Working | -1.1573966776483 | 1 |
| Terrier - Working | -2.10163224760861 | 1 |
| Village_Dog - Working | -3.23679884447629 | 0.0924721182364208 |
| Wolf - Working | 2.64939879295282 | 0.616858664203632 |
| Wolf_Antwerpen - Working | 1.54956746704426 | 1 |
| Wolf_XIX - Working | 1.52993197643757 | 1 |

SI Table 8: **Pairwise Comparison of rECV (regression residuals) between Neolithic and modern wolves and dogs' groups** using Dunn's test with Bonferroni correction. Z values and adjusted p-values are reported. *: p values statistically significant.
